## Supporting Information for "AvianLexiconAtlas: A database of descriptive categories of English-language bird names around the world"

**S1 Appendix. AvianLexiconAtlas Database Files.** The data, glossary, and gazetteer reported in this article can be accessed at https://github.com/ajshultz/AvianLexiconAtlas.

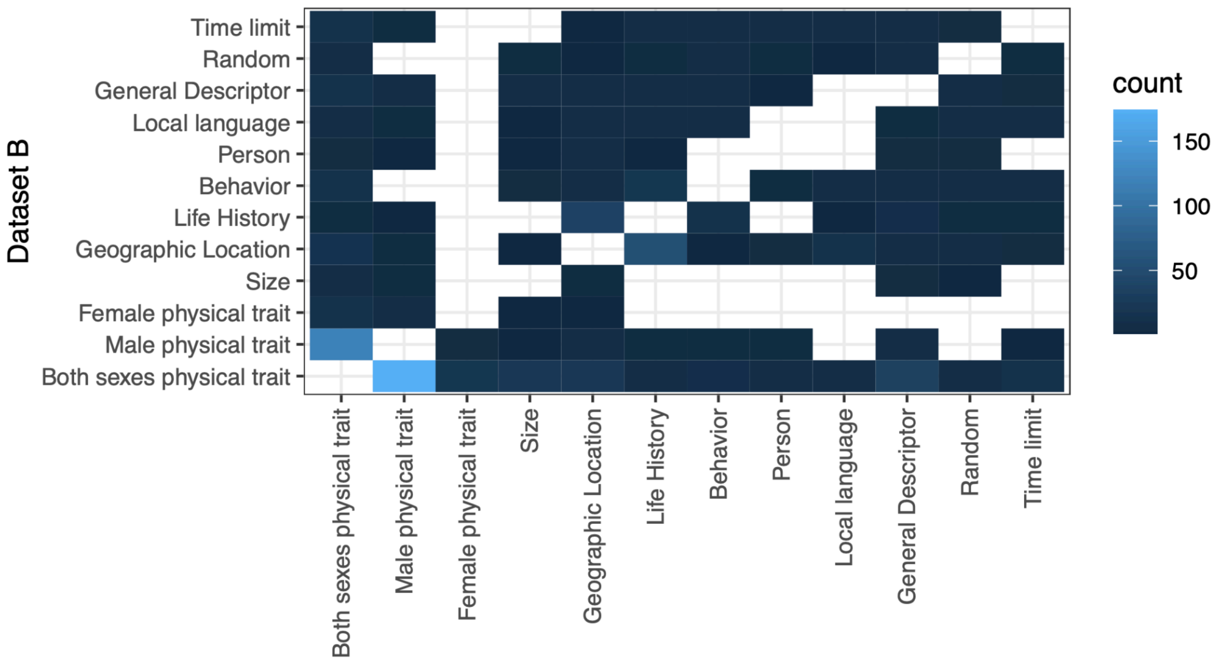


**S1 Fig. Summary of mismatched category assignments in the 915 species common names assigned to different categories in Dataset A and Dataset B.**

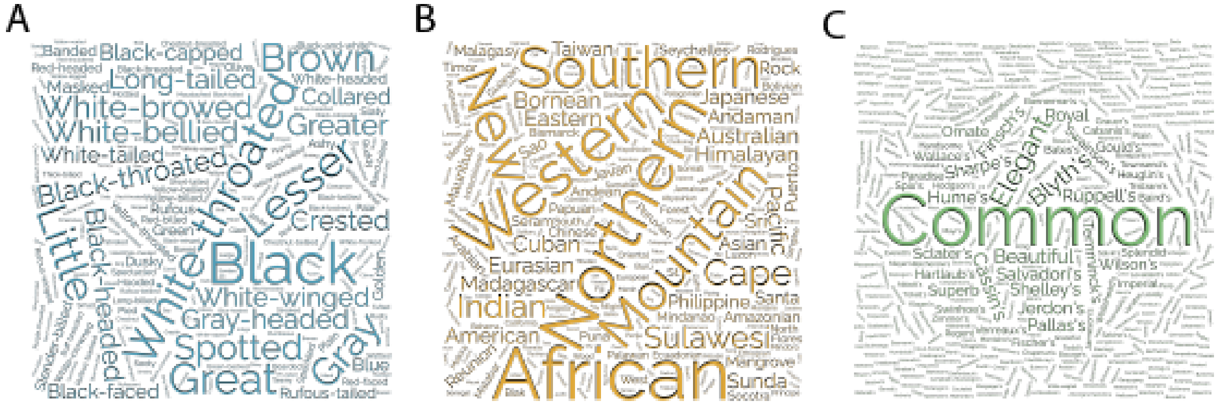


**S2 Fig. Word cloud frequencies of terminology in common names.** Word cloud frequencies of (A) avian physical traits, (B) avian natural history traits, and (C) human-centered terminology. Names are scaled according to frequencies in each dataset.

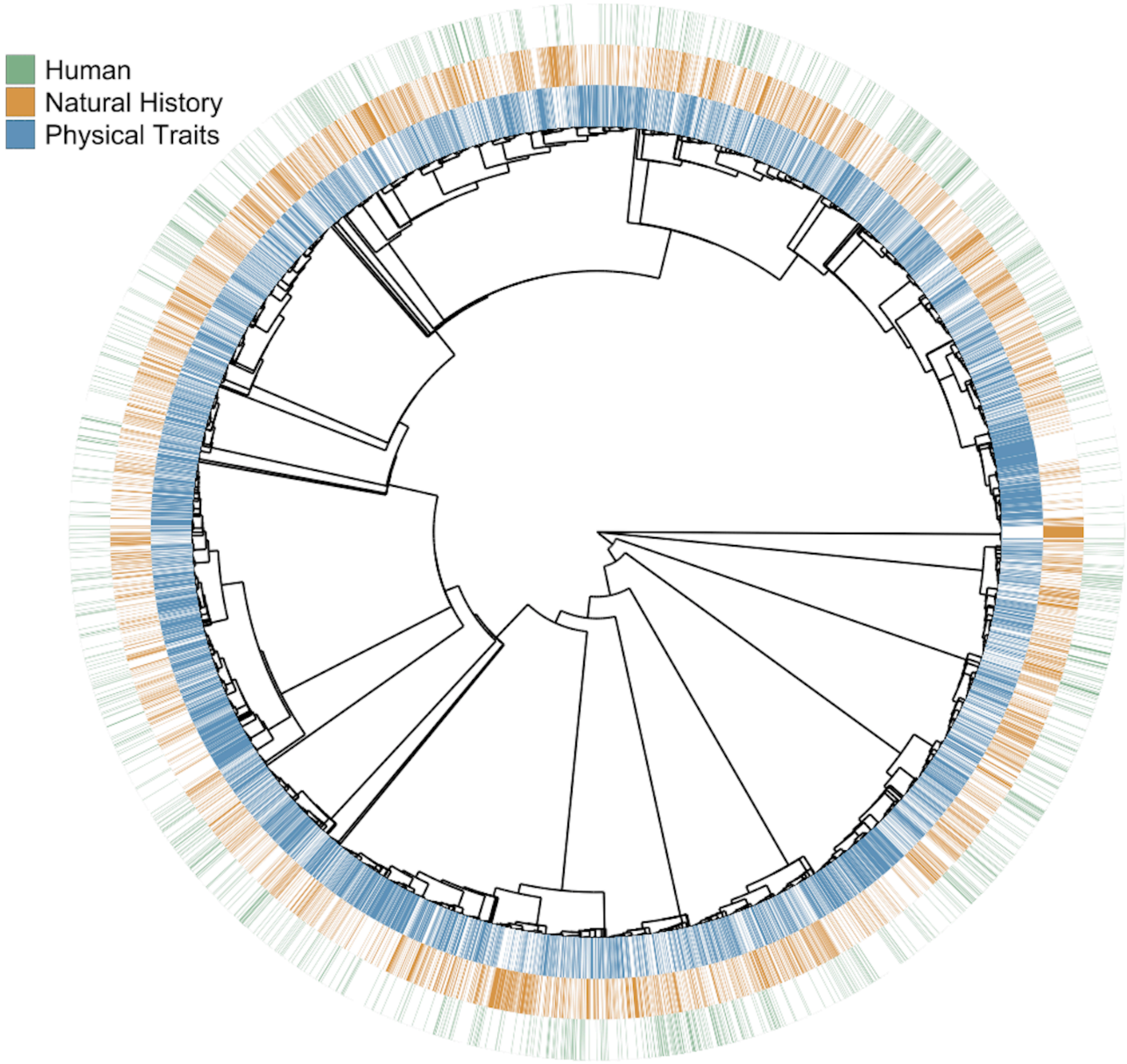


**S3 Fig**. **Cladogram of the categories assigned to the English common names of 10,775 species of birds.** Categories of the English common names identified at the tips of the branches, based on color. Cladogram adapted from [38]. The inner circle includes species names associated with the general category of avian physical traits (both sexes physical trait, male physical trait, female physical trait, size). The middle circle includes species names associated with the general category of avian natural history (behavior, geographic range, natural history), and the outer circle includes names associated with the general category of human-centered terminology unrelated to the biology of the species. See Table 1 for detailed explanations of each category.

**S1 Table. Calculation of D statistics for the phylogenetic structure of categories.** Results of Fritz & Pervis' [40] *D* statistic calculations for the phylogenetic structure of each of the grouped categories: physical traits, natural history, and human-constructed terminology. For each grouped category, species assigned to the category were represented by a state of 1 and the remaining species assigned to other categories were represented by a state of 0. *D* is calculated by scaling the observed sum of sister-clade differences, $\Sigma d_{obs}$, with the mean values of the sum of sister-clade differences for 1,000 simulated trait distributions on the tips of the same phylogeny based on randomly reshuffling the trait values, $\Sigma d_r$, and trait evolution under Brownian motion $\Sigma d_b$: $D = [\Sigma d_{obs} - mean(\Sigma d_b)/[mean(\Sigma d_r) - mean(\Sigma d_b)]$. An estimated *D* close to 1 represents a random distribution of a binary trait among related species on the phylogeny, while an estimated *D* close to 0 represents a clumped distribution of a binary trait among related species that would be expected under the Brownian motion model of evolution. Calculations were completed using the R package *caper* 1.0.3 [41].

| | **Physical Traits** | **Natural History** | **Human Terminology** |
|---|---|---|---|
| **Count of 0 states (no category assigned)** | 4565 | 7361 | 9624 |
| **Count of 1 states (category assigned)** | 6210 | 3414 | 1151 |
| **Observed sums of sister-clade differences,** $\Sigma d_{obs}$ | 3757.153 | 3415.479 | 1761.211 |
| **Mean sums of sister-clade differences of random reshuffling of traits,** $mean(\Sigma d_r)$ | 4560.358 | 4087.216 | 1906.918 |
| **Mean sums of sister-clade differences of trait distributions under Brownian motion,** $mean(\Sigma d_b)$ | 1595.227 | 1451.21 | 733.921 |
| **Estimated *D*** | 0.729 | 0.745 | 0.875 |
| **Probability *D* significantly different from 1 (no phylogenetic structure)** | < 0.001 | < 0.001 | < 0.001 |
| **Probability *D* significantly different from 0 (Brownian motion)** | < 0.001 | < 0.001 | < 0.001 |
